# Tachykinin neuropeptides are involved in axonal and synaptic differentiation of the pioneer motor axon in zebrafish

**DOI:** 10.64898/2026.06.24.734198

**Authors:** Sofiia Ushakova, Daniela Zöller, Anja Bretschneider, Thomas Becker, Catherina G. Becker, Ana-Maria Oprişoreanu

## Abstract

In contrast to many other developing systems, in which axon pathfinding and synaptogenesis are separated in time, the pioneering axon of the individually identifiable caudal primary motor neuron in embryonic zebrafish forms *en passant* synapses during its stereotypical ventral growth. How simultaneous synaptic differentiation and axon pathfinding are coordinated is not fully understood. Here we ask what the role of the *tac1* gene, coding for the synaptic tachykinin neuropeptides, is in this unique axon differentiation process. The gene is expressed during axon outgrowth and its disruption results in increased branch length of CaP axons and subtle morphological defects of the pre-synapse. These abnormalities are accompanied by a robust ∼1.5-fold increase in motor neuron activity and in spontaneous early contractions in *tac1*-deficient embryos. Furthermore, pharmacological inhibition of the tachykinin receptor (Tacr1) leads to altered CaP axonal morphology, mimicking the axonal phenotype observed in *tac1*-deficient zebrafish. These findings suggest that tachykinin neuropeptides modulate formation and activity of *en passant* synapses and prevent aberrant axon branching during growth of zebrafish motor axons.

**HIGHLIGHTS:**

- *tac1* refines CaP primary motor axon development in zebrafish
- Loss of *tac1* disrupts presynaptic maturation at the horizontal myoseptum
- *tac1* mutants show elevated motor neuron activity and spontaneous contractions

## INTRODUCTION

Motor neuron development in zebrafish (*Danio rerio*) is ideally suited to the analysis of growth and differentiation of individually identifiable pioneer neurons *in vivo.* Consequently, this process has been extensively investigated, showing that motor axon differentiation is tightly regulated by various genes (Beattie, 2001; Drapeau et al., 2002). Based on their morphology and differentiation time, motor neurons are classified into primary motor neurons (PMNs) and secondary motor neurons (SMNs) (Myers et al., 1986). Four distinct primary motor neurons have been identified: caudal primary (CaP), middle primary (MiP), rostral primary (RoP) and variable primary (VaP) neurons(Beattie et al., 2002; Myers et al., 1986). Primary motor neurons begin to differentiate at around 9 post-fertilization (hpf), and by 16 hpf their axons exist the spinal cord ventrally. The axons follow a common mid-segmental path until they reach the horizontal myoseptum (HM), an intermediate target, where they pause and form synapses with the muscle pioneer cells. Subsequently, the axons’ paths diverge, with the CaP axon growing ventrally, the MiP axon dorsally and the RoP axon laterally along the trunk. Secondary motor neurons, which differentiate around 15 hpf, begin extending their axons around 33 hpf and follow the trajectories established by the PMNs (Moreno & Ribera, 2009). Notably, the CaP axon pioneers the common path up to the HM and its individual ventral path beyond the HM.

CaP axon growth is closely linked with the continuous formation of *en passant* synapses, the largest of which is at the HM. In the mammalian CNS, *en passant* synapses have been described in telencephalic glutamatergic projection neurons in mice (Zhu et al., 2021), as well as the adult visual cortex (Stettler et al., 2006). At the muscle level, *en passant* synapses are only transiently present during neuromuscular development (Sheard and Duxson, 1997). In *Caenorhabditis elegans* and in the olfactory bulbs of the common garter snake *Thamnophis sirtalis, en passant* synapses persist throughout the organism’s lifetime (Lanuza and Halpern, 1998; Mizumoto and Shen, 2013).

Increasing evidence suggests that axon growth and synapse formation of zebrafish motor axons are tightly coordinated during development. For example, the receptor tyrosine kinase MuSK not only controls postsynaptic acetylcholine receptor pre-patterning but also regulates axon growth cone guidance and synapse localization, indicating that common molecular mechanisms can coordinate both structural and functional maturation of the developing neuromuscular system (Jing et al., 2009, 2010).

A previous screen for differentiation genes associated with CaP axon outgrowth indicated chondrolectin (*chodl*) is one of the genes expressed in motor neurons and regulates synapse-dependent axon growth, exerting its function through a direct interaction with collagen XIX (ColXIX) deposited at the HM. In the absence of *chodl*, CaP axons stall at the HM and synaptogenesis is impaired (Oprişoreanu et al., 2019; Zhong et al., 2012).

In addition to *chodl*, tachykinin precursor 1 (*tac1*) has been shown to be strongly expressed during CaP axon outgrowth, but its function during that early developmental phase is unknown (Zhong et al., 2012). *tac1* encodes neuropeptides that function as neurotransmitters and neuromodulators in the central and peripheral nervous systems. Substance P and neurokinin A (NKA) (Ogawa et al., 2012) are derived from *tac1*. Tachykinins (*tac1*, *tac3*, *tac4*) are widely expressed throughout the central nervous system, where they regulate pain and stress responses, and modulate neuroinflammation (Nässel et al., 2019). In mammals, Tac1-expressing neurons have been implicated in long-range projections that modulate axonal circuitry (He et al., 2024; Hennessy et al., 2017), while studies in larval zebrafish (4 days post-fertilization, dpf) suggest a role for *tac1* in regulating synaptic transmission and motor behaviour (Dill et al., 2024).

Here we find that full deletion of the coding sequence of *tac1* and pharmacological receptor perturbation alters motor axon arborization and morphology of the pre-synapse, leading to increased calcium-dependent neuronal activity and elevated spontaneous movements of embryos. Hence, *tac1* expression is necessary for correct CaP pioneer axon growth and *en passant* synapse formation.

## RESULTS

### tac1 somatic mutants display aberrant CaP axonal phenotype

In a previous expression screen aimed at identifying genes involved in motor axon differentiation, inhibition of LIM homeodomain (LIM-HD) transcription factors resulted in motor axons failing to exit the spinal cord. Two genes coding for neuropeptides, calcitonin-related polypeptide alpha (*calca*) and tachykinin precursor 1 (*tac1*), were found to be highly expressed in spinal motor neurons during axon growth (Zhong et al., 2012). To investigate the potential impact of these genes on motor axon differentiation, we used a functional screening approach (Keatinge et al., 2021) in which we generated somatic mutants for both genes in the *mnx1:GFP* transgenic line, where GFP expression under the *mnx1* promoter allows visualization of CaP axons (Flanagan-Steet et al., 2005).

Gene targeting was achieved by injecting highly active CRISPR RNAs (haCRs) into fertilized eggs, followed by restriction fragment polymorphism (RFLP) analysis (Keatinge et al., 2021), which showed >90% effectiveness in disrupting the targeted restriction enzyme recognition sites for both targeted genes (Suppl. Fig. 1 A, D). *tac1* somatic mutants also exhibited an 89% reduction in mRNA level (Suppl. Fig. 1E). Potential side effects from the injections were controlled by injecting control CrRNA in the same line.

At 26 - 28 hpf, CaP axons were analyzed for aberrant growth, including truncation, excessive branching, and/or absence of axons. In *tac1* somatic mutants, there was a two-fold increase in the percentage of aberrant CaP motor neurons relative to control (25.5% control CrRNA vs 51.19% *tac1* CrRNA; Suppl. Fig. 1G, H). Specifically, *tac1* somatic mutants showed a 40% increase in both average axon branch length (Suppl. Fig. 1J) and number of axon branches (Suppl. Fig. 1K), compared to the CrRNA control group. No significant difference was observed in total axon length in *tac1* somatic mutants (Suppl. Fig. 1I). The *calca* gene was effectively disrupted, as shown by RFLP, but this did not affect axonal outgrowth, compared to control group (Suppl. Fig. 1B, C), such that *calca* disruption only served as an additional negative control in this functional screen. Hence our screening approach indicated potential roles of *tac1* in CaP primary motor axon that warranted further investigation.

### tac1 *mRNA* expression in spinal motor neurons is axon outgrowth-dependent

As a first step to analyze the role of *tac1*, we investigated the *in vivo* presence of *tac1* in spinal motor neurons by RNA fluorescence *in situ* hybridization (HCR RNA-FISH) in *mnx1:gfp^+^* embryos comprising CaP motor axon ventral growth. At 22 hpf, *tac1* mRNA signal was detected in the somata of *mnx1:gfp^+^* spinal motor neurons, but not in other spinal cells. (Fig. 1A). Consistent with previous findings (Zhong et al., 2012), *tac1* expression was detected in spinal cord segments, in which CaP axons had just exited the spinal cord and in those in which the axons had grown beyond the HM (Fig.1A, Region 1 - Box 1, Region 2 - Box 2.1), but not in the most caudal, developmentally younger region, where *mnx1:gfp^+^* neurons were present, but axons had not visibly exited the spinal cord (Fig. 1A, Region 2 - Box 2.2). Hence, expression of *tac1* roughly coincided with CaP axons exiting the spinal cord and with ventral growth along the mid-segmental pathway beyond the horizontal myoseptum.

**Figure 1:**
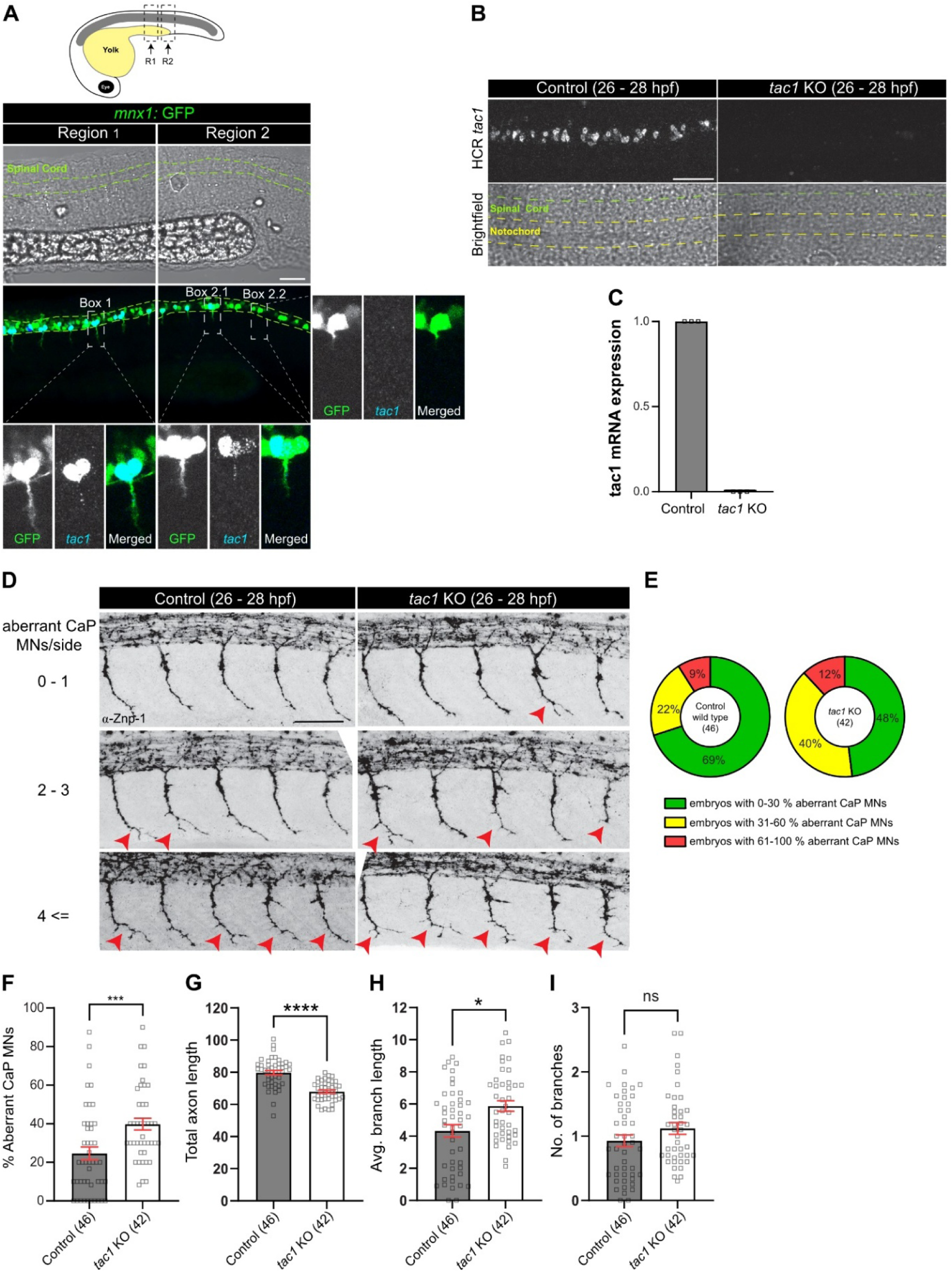
Loss of *tac1* coding sequence alters CaP axonal morphology. **(A)** Lateral trunk views of the 22 hpf (B) *mnx1*:GFP transgenic embryos. CaP motor neurons, whose axons have exited the spinal cord, show *tac1* expression in the cell body (Box 1). This includes the CaP axons that have just exited the spinal cord (Box 2.1). The more caudal, developmentally younger CaP motor neurons (no axon growth detected, Region 2) do not show any detectable *tac1* expression (Box 2.2). **(B)** *tac1* gene expression is no longer detected in the spinal cord of 26 - 28 hpf *tac1* KO embryos compared to control. **(C)** *tac1* KO have undetectable levels of *tac1* mRNA, as assessed by qRT-PCR. **(D)** Lateral trunk views of the 26 - 28 hpf *tac1* KO and control embryos. CaP axons are visualized using anti-Znp-1 antibodies. Red arrows indicate aberrant CaP axons. **(E)** Distribution in percentage of the CaP axonal phenotype per experimental group. Green area - 0 and 30% aberrant CaP motor neurons, yellow area - 31 and 60%, and red area - 61 and 100%. **(F)** Quantification of the percentage of aberrant CaP motor neurons (Mann-Whitney test, ***p = 0.0004). Analysis of the total axon length **(G)** (unpaired t-test, ****p < 0.0001), average branch length **(H)** (Mann-Whitney test, *p = 0.0127) and number of branches **(I)**. Error bars show SEM. Scale bar: 50 µm

### Loss of *tac1* results in aberrant CaP axonal phenotype

To confirm the functional role of *tac1*, we generated a stable germline loss-of-function mutant by removing the entire *tac1* open reading frame (ORF) using two CrRNA guides and CRSPR/Cas9 (Suppl. Fig. 2A). Following genotyping (Suppl. Fig. 2A), a germline stable *tac1* knockout line (*tac1* KO) was raised. qRT-PCR (Fig. 1C) and HCR RNA-FISH at 26 - 28 hpf (Fig. 1B) confirmed complete absence of *tac1* transcript in the mutant. Gross developmental assessment of *tac1* KO embryos, including eye diameter measurements, revealed no significant abnormalities at 26 - 28 hpf. However, a small reduction in body length was observed (10% decrease, Suppl. Fig. 2B). *tac1* KO embryos and larvae were viable and developed normally into adulthood.

Analysis of the *tac1* KO CaP axonal phenotype was performed at 26 - 28 hpf using anti-Znp-1 antibodies to label the presynaptic compartments (pre-synaptic marker, Synaptotagmin-2) along the extending motor axons and their branches. The analysis revealed a 1.61-fold increase in the percentage of aberrant CaP axons compared to the non-homozygous control group (24.62% control vs 39.83% *tac1* KO) (Fig. 1F). To further characterize the morphology of CaP axons, we measured total axon length, as well as the number and length of branches. Total axon length was significantly reduced by 15% in *tac1* KO compared with wild-type embryos (Fig. 1G). Additionally, the average branch length was increased by approximately 35% in the *tac1* KO group (Fig. 1H), consistent with observations in *tac1* somatic mutants (Suppl. Fig. 1J). No statistically significant difference was observed in the number of branches between the groups (Fig. 1I). These observations confirmed that *tac1* expression was necessary for correct outgrowth of CaP axons.

### Pharmacological block of the Tacr1 receptor induces aberrant CaP axonal differentiation

To test whether *tac1* products likely acts via the Tacr1 receptor, we used a pharmacological approach to block it. The products of the *tac1* gene, substance P (SP) and neurokinin A (NKA), bind to G-protein-coupled tachykinin receptors. In zebrafish, two genes, *tacr1a* and *tacr1b,* encode two subtypes of the Tacr1 receptor: Tacr1a and Tacr1b. The *tacr1a* receptor is expressed in various regions of the brain, while *tacr1b* is expressed in the same regions as *tacr1a* but is additionally found in the spinal cord (Lopez-Bellido et al., 2013), and at lower levels in muscle tissue (Sur et al., 2023).

The Tacr1 receptor was pharmacologically blocked using Rolapitant, a selective and competitive antagonist of human SP/NKA receptor NK1, which resulted in a 2.13-fold increase in the percentage of aberrant CaP MNs at 26 - 28 hpf compared to vehicle-treated control (27.55% DMSO-control vs 58.89% Rolapitant) (Fig. 2A, B). Further axonal analysis revealed that Rolapitant treatment caused a 9.69% reduction in total axon length, a 23.62% increase in average branch length, and a 44.5% increase in the number of branches compared to DMSO-treated embryos at 26 - 28 hpf (Fig. 2D, E, F). These results closely resemble the phenotype observed in the *tac1* somatic mutants and KO animals. These findings indicate that the pharmacological inhibition of Tacr1 receptor disrupts normal CaP axon development, highlighting the importance of Tacr1-mediated signalling in motor axon growth and branching.

**Figure 2:**
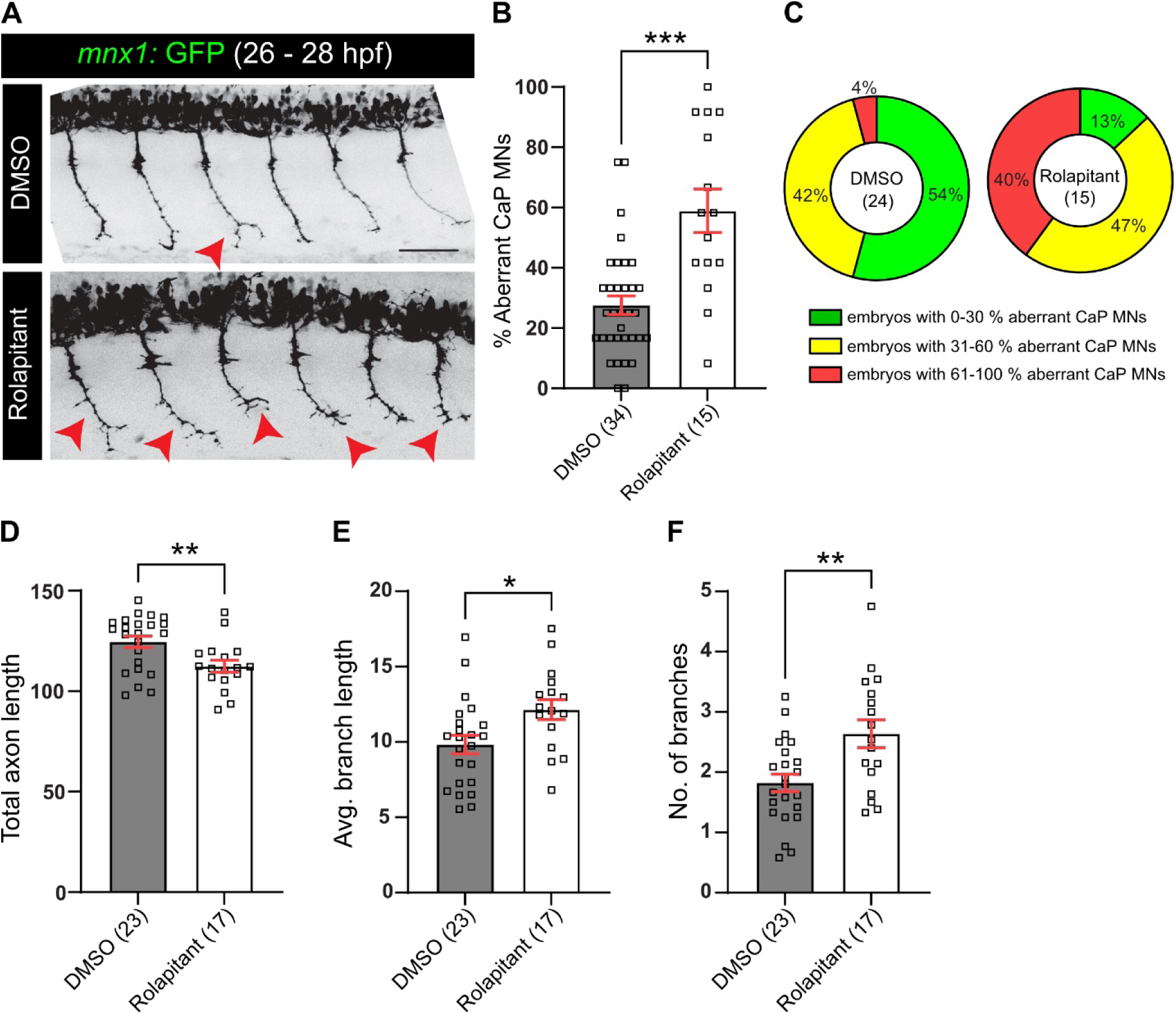
Tacr1 receptor inhibition leads to altered CaP axonal morphology. **(A)** Lateral trunk views of the 26 – 28 hpf *mnx1*:GFP embryos incubated with Tacr1 receptor inhibitor, Rolapitant. Red arrows indicate aberrant CaP axons. **(B)** Quantification of the percentage of aberrant CaP motor neurons (Mann-Whitney test, DMSO vs Rolapitant, ***p = 0.0001). **(C)** Distribution in percentage of the CaP axonal phenotype per experimental group. Green area - 0 and 30% aberrant CaP motor neurons, yellow area - 31 and 60%, and red area - 61 and 100%. Analysis of total axon length (**D**, unpaired t-test, **p = 0.0063), average branch length (**E**, unpaired t-test test, *p = 0.0144), andnumber of branches (**F**, unpaired t-test, **p = 0.0060). Error bars show SEM. Scale bar: 50 µm.

### *tac1* KO embryos exhibited morphological presynaptic defects

The loss of *tac1* could affect axon growth directly or as a secondary consequence of synaptic aberrations at the HM and other synaptic sites. We therefore decided to analyze motor axon synapses with the muscle tissue in detail. The horizontal myoseptum (HM) is a major synaptic site, where CaP motor neurons form *en passant* synapses (Beattie et al., 2002; Oprişoreanu et al., 2019). Hence, the absence of *tac1* gene could disrupt axonal branching at the synaptic sites, which serve as starting points for axon branching (Javaherian and Cline, 2005; Panzer et al., 2005).

To investigate potential changes in synaptogenesis of CaP motor neurons at the HM, wild-type control and *tac1* KO zebrafish embryos were immunolabeled at 26 - 28 hpf using Znp-1 antibodies and AChR antibodies to label the pre-and postsynaptic compartments (Fig. 3A). We quantified the total area of the pre-and post-synaptic regions, their overlap, the number of discernible synaptic labelling areas (cluster number), and the intensity of the labelling in high-resolution microscopy (Oprişoreanu et al., 2019). The total area of the pre-, post-and overlapping synaptic regions was unchanged between control and *tac1* KO embryos (Fig. 3B). The number of synaptic clusters in the presynaptic compartment and the overlapping synaptic regions decreased by 9% and 11%, respectively, in *tac1* KO embryos. The labelling intensity of Znp-1 was increased by 13% in *tac1* KO embryos, while the mean intensity of AChR labelling remained unchanged (Fig. 3B). These findings indicated that lack of *tac1* gene induced a subtle increase in the mean intensity of the presynaptic marker and a decrease in the number of discernible presynaptic and synaptic puncta, which could potentially influence neuronal activity.

**Figure 3:**
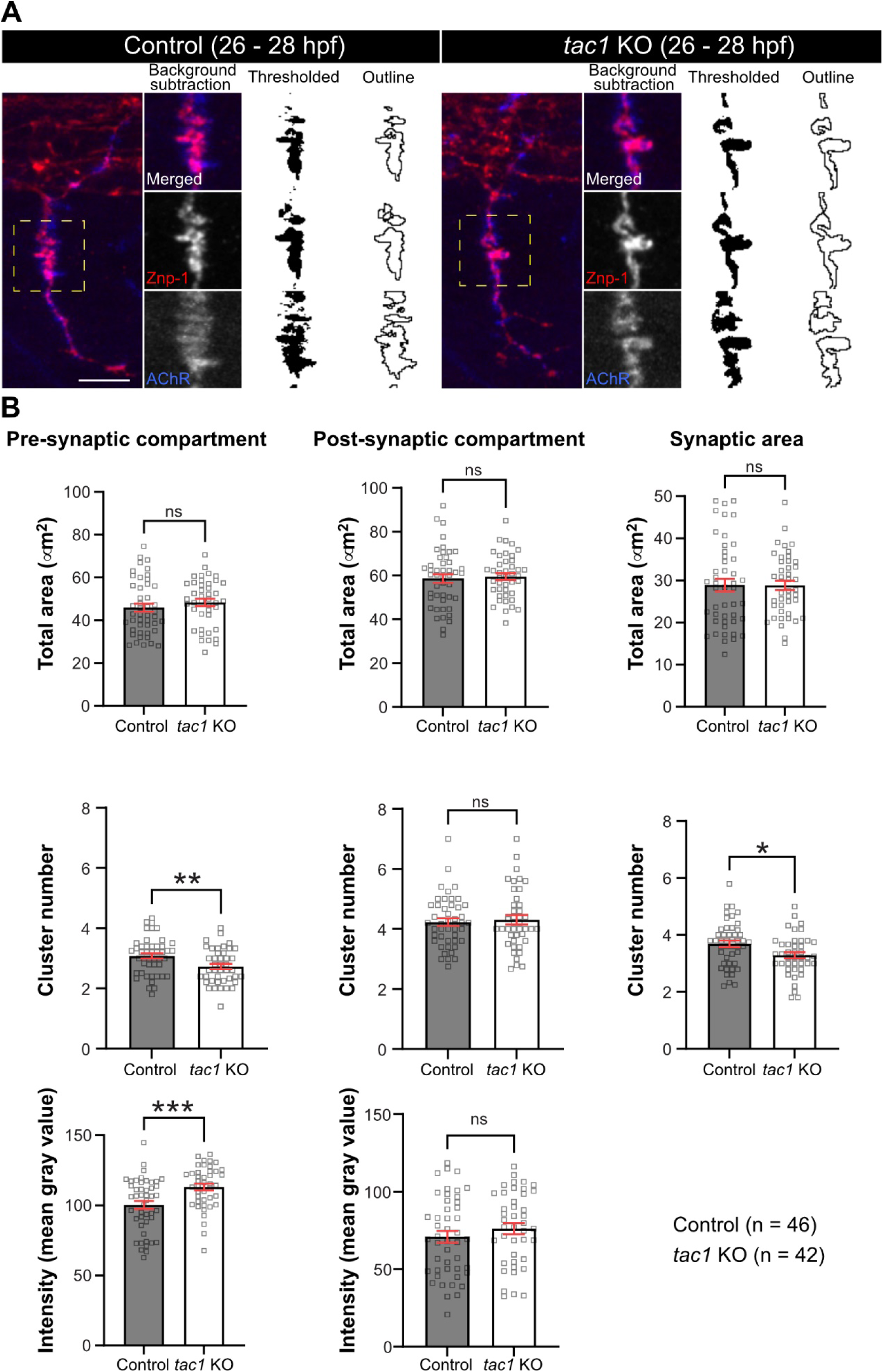
Loss of *tac1* gene induces defects in synapse morphology. **(A)** Representative images of CaP motor neurons. Pre-synapse is labelled by anti-Znp-1 antibodies, and post-synapse by anti-AChR antibodies. **(B)** Analysis of the synapse at the horizontal myoseptum (marked by the yellow dashed box in A). Total area of the pre-and post-synapse is unchanged between *tac1* KO and control groups. *tac1* KO have fewer pre-synaptic and synaptic clusters compared to control (pre-synapse - unpaired t-test, **p = 0.0076; synaptic area - unpaired t-test, *p = 0.0136). Intensity of Znp-1 labelling is increased in *tac1* KO embryos compared to control (unpaired t-test, ***p = 0.0010). Scale bar: 20 µm. Error bars show SEM.

### *Tac1* KO embryos showed increased motor neuron activity

To determine whether motor neuron activity of spinal neurons was indeed altered by disruption of *tac1*, the CaMPARI2 sensor plasmid was used. The CaMPARI2 protein is photoconverted from green to red in the presence of UV light depending on intracellular calcium levels and thus provides a cumulative read-out for neuronal activity/intracellular calcium levels (Moeyaert et al., 2018). We acutely injected a FLAG-tagged CaMPARI2 plasmid and found expression of the protein under a pan-neuronal promoter (HuC) in motor neurons (identified by their ventral position, large size and axon exiting the spinal cord) and other spinal neurons.

To detect activity of these neurons, we illuminated 26 - 28 hpf embryos for 10 minutes with UV light and determined the ratio of photoconverted protein (anti-EosFP Red antibody) to total protein expression (anti-FLAG antibody). This indicated an 52% increase in relative fluorescence intensity in motor neurons of *tac1* KO embryos, compared to wildtype embryos. No significant change in CaMPARI2 conversion was observed in non-motor neurons (Fig. 4B, C). The result indicates selectively increased neuronal activity in motor neurons in the absence of *tac1* expression, and is in line with increased presynaptic marker intensity in motor axons of *tac1* KO embryos.

**Figure 4:**
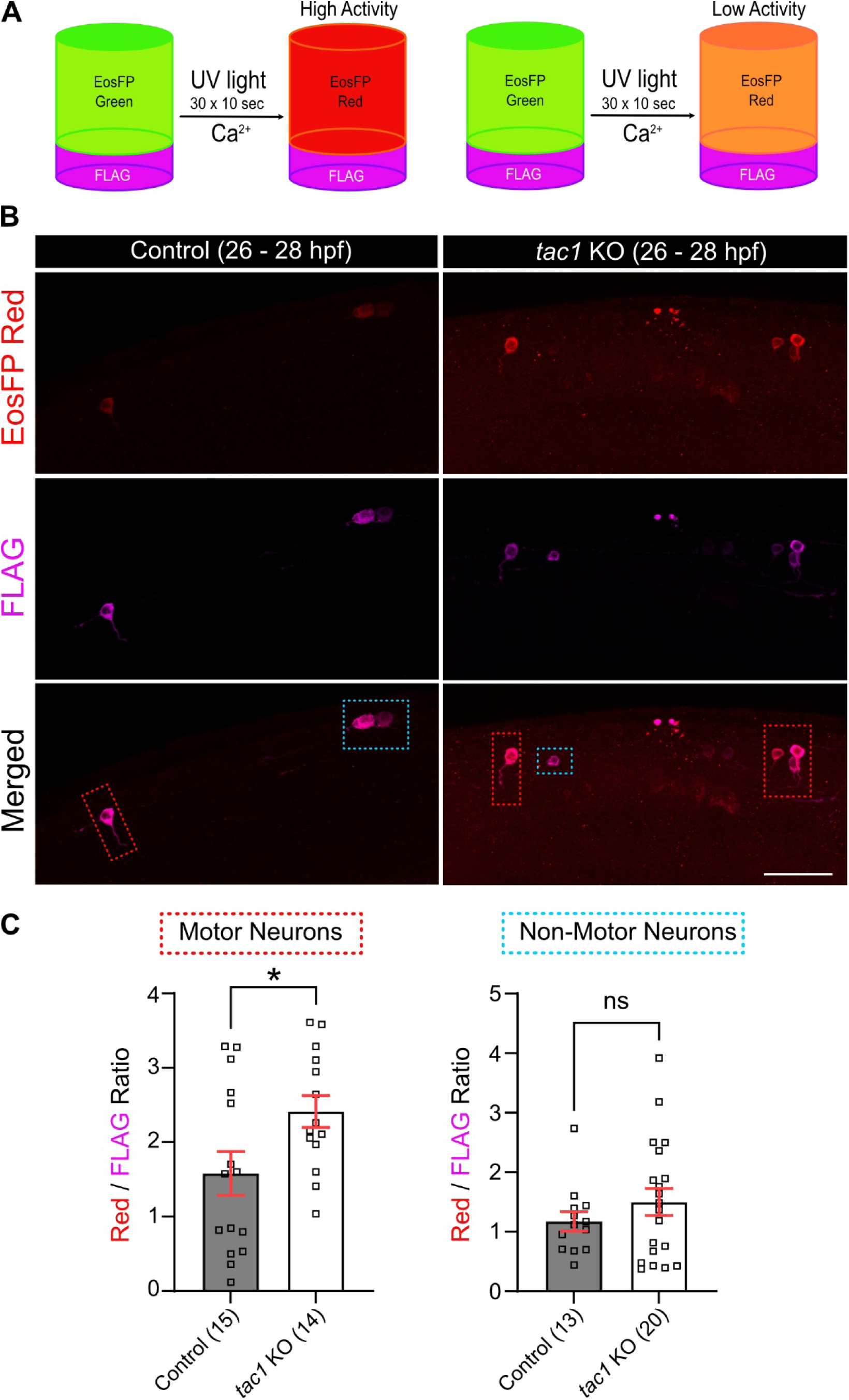
Motor neuron activity is increased in *tac1* KO embryos. **(A)** Pictogram depicting the CaMPARI2 sensor conversion. **(B)** Representative images of motor neurons overexpressing the CaMPARI2 sensor. Anti-FLAG antibodies labels both photoconverted and non-photoconverted construct. Motor neurons are marked by the red dotted boxes, while the non-motor neurons by the turquoise boxes. **(C)** Quantification of EosFP Red / FLAG signal ratios (Motor neurons, unpaired t-test, *p = 0.0318). Scale bar: 50 µm. Error bars show SEM.

### The frequency of spontaneous early contractions was increased in *tac1* KO embryos

Based on the observed synaptic defects and increased calcium-dependent neuronal activity of motor neurons, we investigated whether muscle activity was also increased as a consequence. Zebrafish embryos show early contractions that are elicited by spontaneous activity of neuro-muscular junctions (Drapeau et al., 2002; Saint-Amant and Drapeau, 1998) Therefore, we tested whether the frequency of these contractions was altered in the absence of *tac1* gene. The earliest motor axon - muscle interaction occur at 18 - 19 hpf, before the CaP axons reach the HM (Myers et al., 1986; Saint-Amant and Drapeau, 1998). For the experimental setup, embryos were maintained under constant ambient temperature and light conditions. Individual embryos were recorded one at a time, alternating between genotypes. At this developmental stage, no difference in coiling behaviour frequency was observed between control and *tac1* KO embryos (Fig. 5A). However, at 24 - 26 hpf, when CaP axons had formed synapses at the HM and extended beyond this choice point, *tac1* KO embryos exhibited a 53% increase in the frequency of spontaneous contractions compared to controls (Fig. 5B).

**Figure 5:**
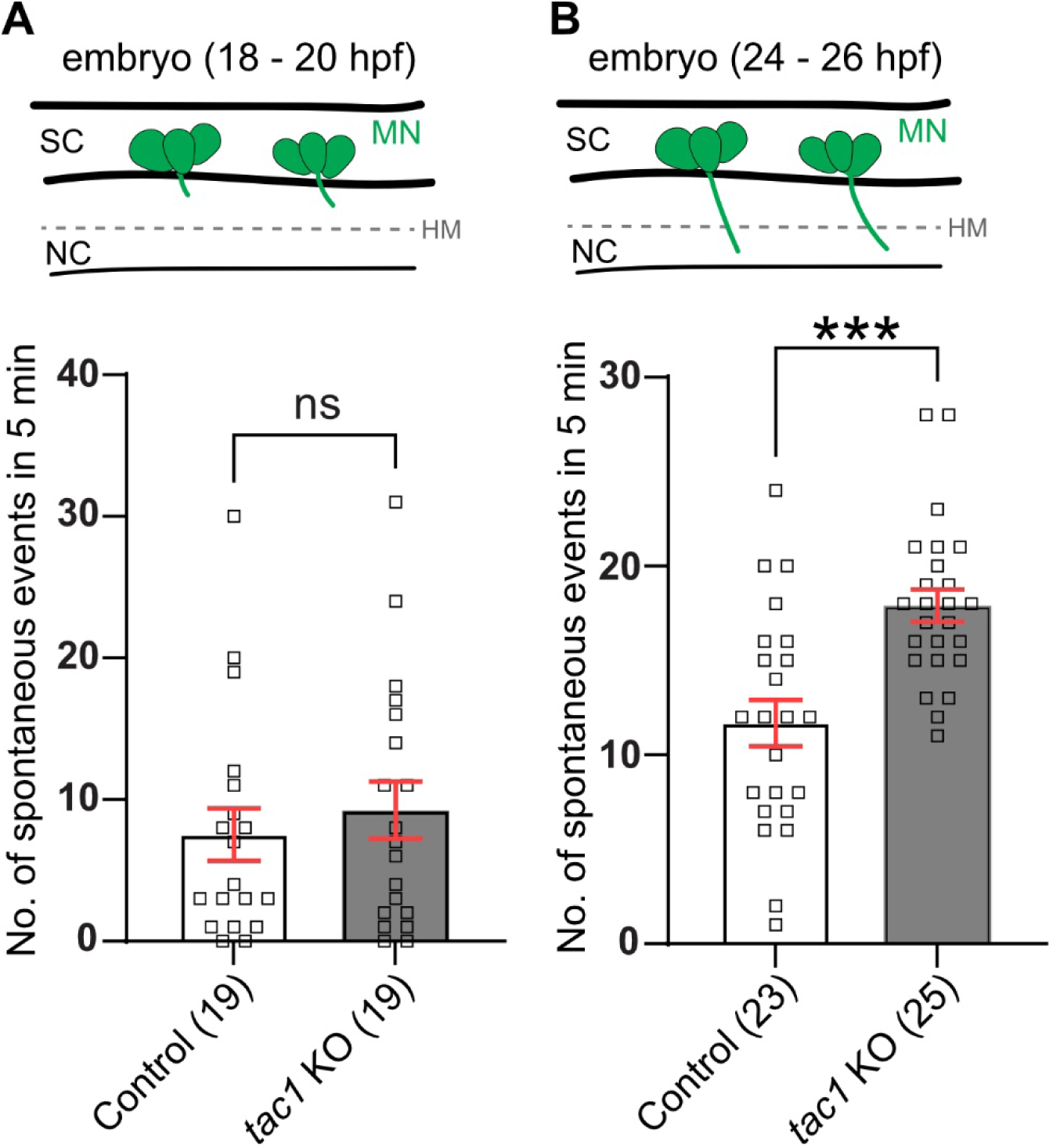
Spontaneous early contractions are increased in frequency in the absence of the *tac1* gene. **(A)** No change in spontaneous contractions is observed in *tac1* KO compared to control at earlier time points in development (18 - 20 hpf). **(B)** 26 - 28 hpf *tac1* KO embryos show more spontaneous contractions events that control group (unpaired t-test, ***p = 0.0001). Error bars show SEM.

In summary, disruption of *tac1* resulted in altered presynaptic organization, increased motor neuronal activity, and elevated spontaneous motor behavior. The increased frequency of spontaneous early contractions indicates altered output of the developing motor system, whereas the changes in synaptic morphology and neuronal activity point to broader disturbances in motor circuit maturation. Together, these findings support a role for *tac1* in coordinating structural and functional aspects of early motor neuron development.

## DISCUSSION

We demonstrate that *tac1* fine-tunes primary motor axon development and early motor system activity in zebrafish. By combining complementary approaches (CRISPR/Cas9 - somatic mutagenesis, stable germline knockout generation, receptor pharmacological blockade, synapse morphology characterization, and neuronal activity measurements) we provide evidence that *tac1* gene disruption alters CaP axonal morphology, affects pre-synapse morphology, increases motor neuronal activity, and elevates spontaneous motor behavior. Together, our findings suggest that *tac1* contributes to the coordination of axon outgrowth and synapse maturation during early motor circuit formation.

### *tac1* regulates motor axon outgrowth

Our somatic CRISPR/Cas9 screen identified *tac1*, but not *calca*, as a candidate gene regulator of CaP axon development. Somatic disruption of *tac1* led to excessive CaP axonal branching, a phenotype that was recapitulated in the stable *tac1* knockout line, supporting the idea that *tac1* is required for stereotypical motor axon outgrowth.

The temporal and spatial expression pattern of *tac1* further supports a developmental role during motor axon extension. *tac1* expression was detected specifically in spinal segments in which CaP axons had exited the spinal cord and extended along their ventral trajectory, whereas expression was absent from younger caudal segments where motor neurons were present, but axon emergence from the spinal cord had not yet occurred. These observations suggest that *tac1* is associated with axon outgrowth rather than early motor neuron development. Previous studies have similarly shown developmentally regulated *tac1* expression in the zebrafish nervous system (López-Bellido et al., 2013).

An interesting possibility is that *tac1* acts within a developmental program controlling sequential stages of motor neuron maturation. Previous work identified *chodl* as an important regulator of motor axon differentiation (Oprişoreanu et al., 2019), while *calca* and *tac1* expression appear later in motor neuron development (Zhong et al., 2012). Our observations raise the possibility that *tac1* functions during an intermediate developmental window, following early differentiation programs (*chodl*) and preceding later maturation steps (*calca*). This timing may also explain why disruption of *calca* in our CRISPR/Cas9 screen at 26 - 28 hpf did not result in a detectable aberrant CaP axonal phenotype. Our previous study had shown that *calca* expression at 22 hpf is detected in the more rostral spinal segments, where motor axons have already extended considerably, whereas *chodl* and *tac1* are abundantly expressed in all segments during earlier stages of motor axon outgrowth (Zhong et al., 2012).

### *tac1* regulates synapse maturation

Our data suggests that *tac1* contributes to presynaptic maturation of the CaP axon at the horizontal myoseptum (HM). In the absence of *tac1*, presynaptic morphological alterations were observed, including reduced numbers of presynaptic and synaptic puncta together with increased labelling intensity of the presynaptic marker Znp-1. Interestingly, these alterations partially resemble defects previously described in *chodl* mutants (Oprişoreanu et al., 2019), raising the possibility that *tac1* provides an additional regulatory layer controlling synapse development.

One possible interpretation of our data is that *tac1* normally helps maintain developing synapses in a relatively immature state during periods of active axon growth. Coordination between motor axon growth and synapse development has been demonstrated for other neuromuscular signaling pathways in zebrafish. For example, Agrin is required for motor axon outgrowth and branching (Kim et al., 2007), whereas MuSK signaling regulates both axon growth cone guidance and synaptic patterning (Jing et al., 2009; Lefebvre et al., 2007). These studies, together with our findings raise the possibility that *tac1* may coordinate motor axon growth and synapse maturation during early motor circuit assembly.

Developmental studies have shown that axon extension and synapse maturation are tightly coordinated processes, with structural plasticity being important during periods of active circuit assembly (Javaherian and Cline, 2005; Panzer et al., 2005). In this context, delayed synaptic stabilization may promote structural plasticity necessary for axon growth and appropriate branching, whereas premature synaptic stabilization, indicated by the observed heightened neuronal activity after *tac1* disruption, may constrain these developmental processes. Thus, loss of *tac1* could alter the balance between axon growth and synapse maturation, leading to abnormal branching and altered presynaptic organization.

The increased Znp-1 signal observed in *tac1* mutants suggests altered organization of the presynaptic terminals. Because Znp-1 marks the synaptic vesicle-associated protein Synaptotagmin-2, the increased signal may reflect an accumulation of synaptic vesicles, altered vesicle trafficking, or increased abundance of SV-associated proteins at the presynaptic terminal. Although the underlying mechanism remains unclear, such alterations could potentially contribute to increased neurotransmitter release and heightened excitability within early motor circuits. Consistent with this interpretation, *tac1* knockout embryos displayed increased motor neuronal activity together with elevated spontaneous motor behavior. Further experimental approaches will be required to determine the mechanism underlying this phenotype.

Importantly, behavioral differences were not observed before CaP axons established synapses at HM, but after synaptic contacts had formed. This temporal relationship suggests that *tac1*-dependent effects become important once developing motor axons transition from axon outgrowth toward synaptic integration and functional circuit assembly. Early spontaneous motor behavior in zebrafish emerge progressively during spinal circuit maturation (Drapeau et al., 2002; Saint-Amant and Drapeau, 1998), supporting the interpretation that altered behavioral reflects disruption of developmental motor circuit refinement. Our findings further extend previous observations describing heightened motor activity in 4 dpf *tac1* mutant zebrafish larvae (Dill et al., 2024). That study established a role for *tac1* in acutely regulating neuronal and behavioral activity at larval stages, and our study reveals an additional developmental role of *tac1* in setting up the motor system.

In mammals, Tac1-derived peptides, Substance P and Neurokinin A, regulate neurotransmission, neuronal excitability, and synaptic signaling (Pennefather et al., 2004; Severini et al., 2002). Together, these findings suggest that tachykinin signaling contributes not only to mature neuronal function but also to structural and functional development of early motor circuits.

### Study limitations and future directions

Several limitations should be considered when interpreting these findings. *tac1* gene encodes two neuropeptides, Substance P and Neurokinin A (Ogawa et al., 2012), and the present experiments do not distinguish which signalling peptide mediates the observed developmental effects. Although pharmacological inhibition of Tacr1 signalling phenocopied major aspects of the mutant phenotype, the cellular localization of the relevant receptors (Tacr1a and Tacr1b) remains unresolved. Limited availability of zebrafish-specific reagents (e.g. antibodies) restricts direct examination of receptor localization during motor neuron development.

In addition, zebrafish possess other tachykinin genes (e.g. *tac3* and *tac4)* (Nässel et al., 2019) whose potential contributions to synapse formation and motor circuit maturation remain unknown. Future studies combining receptor-and peptide-specific manipulations, and cell-type-specific analyses will be important to further define the mechanisms through which tachykinin signalling shapes motor neuron development.

## Conclusion

In conclusion, our findings identify *tac1* as an important modulator of CaP motor axon development in zebrafish. *tac1* influences axonal morphology, presynaptic organization, motor neuronal activity, and motor behaviour of this pioneer axon, indicating a broader role in coordinating structural and functional maturation at the earliest stages of motor circuit assembly.

## MATERIALS AND METHODS

### Animals

All zebrafish lines were raised and kept under standard conditions (Westerfield, 2007). Zebrafish experiments were performed under state of Saxony licenses TVV 35/2025, TV vG 3/2023. For experimental conditions, embryos up to an age of 28 hours post-fertilization (hpf) were used. The following lines were used: wild type (AB) and Tg(*mnx1:*GFP)^ml2tg^, abbreviated as *mnx1*:GFP (Flanagan-Steet et al., 2005).

### Generation of *tac1* and *calca* somatic mutants

Somatic mutations were generated based on adapted protocol from Keatinge et al. (Keatinge et al., 2021). CRSPR gRNAs were designed using the Milipore or IDT CRISPR Design tools (ENSDART00000146947.2/ NM_001256391 reference sequence for *tac1*, ENSDART00000079112.6/ NM_001002471 reference sequence for *calca*). CrRNAs were chosen based on specificity (above 95%), efficiency (above 50), and on the availability of restriction enzyme recognition sites that would be destroyed by an efficient CrRNA. Microinjections were performed in one-cell stage Tg(mnx1:GFP) eggs using the PV 820 Pneumatic PicoPump microinjector by WPI with vacuum eject pressure of 20 psi. Each egg was injected with 1 nL of CrRNA mix containing: 1 µl of Tracer (5 nM, TRACRRNA05N, Sigma-Aldrich), 1 µl of CrRNA (20 µM, Sigma-Aldrich), 1 µL of phenol red (P0290, Sigma-Aldrich) and 1 µL of Cas9 (20 µM, M0386M, New England Biolabs). *tac1* CrRNA1: 5’-CAAGTTTTTGGAGAGGAATT-3’ and *tac1* CrRNA2: 5’-AAGACCTTGATTACTGGAC-3’. *calca* CrRNA1: 5’-AGGCGATGGGTCACGCATG-3’ and *calca* CrRNA2: 5’-CAAAGAATTCATGCAGATGA-3’.

For CrRNA efficiency testing, genomic DNA was extracted from control and CrRNA-injected embryos using 50 mM NaOH, followed by boiling at 95°C for 15 min. Following neutralization with 1 M Tris-HCl, PCR was performed using the following primers: *tac1* (both CrRNA) fw 5’-GGAGCAACATGAAATTTATTTTACC-3’ and rv 5’-CAGTAGGTTTTTTTTGATCTGCAAG-3’, *calca* CrRNA1 fw 5’-TGTTACGACACAGAAGAGAGC-3’ and rv 5’-CTACTCCTGACTGAGCCTC-3’, *calca* CrRNA2: fw 5’-GCCCGCACTGGA ATCGTC-3’ and rv 5’-GTTTTCCTCGGTAGCTTGTTG-3’. CrRNA efficiency was tested by restriction fragment length polymorphism analysis (RFLP) in a proportion of larvae for each experiment (*tac1* CrRNA1 with MluCI, *tac1* CrRNA2 with BsrI, *calca* CrRNA1 with NspI, *calca* CrRNA2 with MslI). All experiments conducted with somatic mutants were carried out with a control injected group with a non-targeting gRNA.

### Generation of *tac1* knockout line

To generate a stable *tac1* knock-out line, the complete *tac1* open reading frame was removed using the following CrRNA guides: AAGAACTGTGTTATATGTGG and TTCCATTCTGTGATACCCTT. The founder selected to generate the stable line had the *tac1* ORF completely removed as assessed by fin-genotyping and sequencing. For genotyping the following primers were used: fw: TTCCTCTACCCGACGCGAAC, rv1: GGTGGAGAGAGCGTGTCACT and rv2: CAGTAGGTTTTTTTTGATCTGCAAG. Fw and rv1 primers were used to detect knock-out animals (693 bp), whereas fw and rv2 were used to identify wild type animals (521 bp). The *tac1* KO morphology was assessed at 28 hpf by measuring the body length and eye diameter using the VAST BioImager (Union Biometrica).

### *tac1 in situ* hybridization chain reaction (HCR)

*tac1* hybridization chain reaction (HCR) was performed using a previously published protocol (Ibarra-García-Padilla et al., 2021). Embryos were fixed for 4h in 4% PFA, followed by over-night storage in methanol at-20 °C. Next day the samples were rehydrated and permeabilized using proteinase K (10 µg/ml in 1xPBST) for 30 min. Hybridization with 3 pmol of *tac1* probe was performed overnight at 37 °C. The following day the HCR amplification was performed using the B3-647 amplifier (Molecular Instruments). Following the HCR amplification, the samples were extensively washed in 5x SSCT and 1x PBST buffer and stored in 70% glycerol. Image was performed using an inverted Zeiss LSM 980 confocal microscope.

### RNA extraction and quantitative RT-PCR

RNA was extracted from 26 - 28 hpf embryos (wild type, *tac1* CrRNA, and *tac1* KO) (50 larvae/group/experiment) using the RNeasy Mini Kit (Qiagen) according to the manufacturer’s instructions. cDNA was synthesized using LunaScript SuperMix (NEB, cat. No. M3010X) according to the manufacturer’s instructions. Reverse transcription - PCR was performed with the following set of primers: β-actin fw 5’-CACTGAGGCTCCCCTGAATCCC-3’ and rv 5’-CGTACAGAGAGAGCACAGCCTGG-3’, *tac1* fw 5’-GTTTTTGGAGAGGAATTGGGTC-3’ and rv 5’-CCGTGTGATCTGTGCATTTG-3’. qPCR was performed using the Luna Universal qPCR Master Mix (NEB, cat. No. M3003L) and the QuantStudio3 machine (Thermo Fisher Scientific). Each condition was normalized to β-actin housekeeping gene and the experimental group (*tac1 CrRNA* or *tac1 KO*) was compared to controls by normalizing to the control group. For statistical analysis, a One-Sample T-test was used.

### Analysis of CaP axonal morphology

To assess CaP axonal morphology, dechorionated Tg(*mnx1*:GFP) 26 - 28 hpf embryos were fixed in 4% PFA for 2 hours at RT. Next, embryos were washed in phosphate buffered saline with 0.1% Tween-20 (PBS-T) 3 times for 5 minutes and stored in 70% glycerol in PBS at 4°C overnight. For imaging, embryos were manually deyolked and mounted in 70% glycerol onto glass slides.

For analysis optically sectioned stacks (z-stack step = 1 µm) of the caudal region (from somite 7) of zebrafish spinal cord were recorded using a Carl Zeiss inverse Imager microscope with an Apotome unit (AxioCam MRm (60N-C) and a LD LC1 Plan-Achromat 20x-objective (NA=0.8)). Images were acquired using the Zeiss Zen Black Software. The z-stacks were converted into maximum intensity projections (MIPs) for each side of the embryo. The percentage of aberrant CaP axons was quantified by analysing 12 - 14 axons per embryo (6 - 7 axons per side) in Fiji following the previously published protocol (Oprişoreanu et al., 2021). CaP axons that were stalled, highly branched or/and missing were categorized as aberrant. To measure total axon length, number of branches and branch length, images were converted to 8-bit format and axons/branches were traced using the Fiji plugin NeuronJ (Meijering et al., 2004). For analysis, the observer was blinded to the experimental conditions.

### Whole-mount immunohistochemistry and synapse quantification

Using a previously published protocol (Oprişoreanu et al., 2019), the 26 - 28 hpf zebrafish *tac1* KO and wild type embryos were manually dechorionated, fixed in 4% PFA and stained for pre-and postsynaptic compartments using the following primary antibodies: mouse Znp-1 (1:100, against synaptotagmin-2, DSHB) and rat mAb35 (1:200, against acetylcholine receptor (AChR) nicotinic alpha 1 subunit, DSHB). Secondary antibodies were Alexa Fluor 594 donkey anti-mouse with pre-adsorption against rat protein (1:400, Jackson ImmunoResearch) and Alexa Fluor 647 anti-rat with pre-adsorption against mouse (1:400, Jackson ImmunoResearch). For analysis of CaP axons in the caudal region (from somite 7) of spinal cord were imaged using an inverted Zeiss LSM 980 confocal microscope (CMCB Light Microscopy Facility of the CMCB Technology Platform, TU Dresden). Synaptic puncta quantification was performed in ImageJ using a previously established protocol (Oprişoreanu et al., 2019). Briefly, a square region of interest was drawn around the HM. For each channel, the background was subtracted, and a 30% threshold was applied to generate a binary image. Using the Analyse Puncta function in Fiji, the outline masks were generated to visualize the synaptic puncta. Total area, puncta number and intensity were analysed for 4 to 5 axons per side (both sides were analysed), and values were averaged per embryo. For quantification the observer was blinded to the experimental condition.

### Neuronal activity using the CaMPARI2 sensor

To measure the neuronal activity, CaMPARI2 sensor was transiently expressed in *tac1* KO and wild type embryos. 25 ng/ml pTol2-HuC(elavl3)-CaMPARI2 (Addgene plasmid no. 137185) was co-injected with 50 ng/ml Tol2 transposase mRNA (Addgene plasmid no. 31831) at the single-cell stage. At 24 - 25 hpf, embryos were screened for the expression of the EosFP Green fluorescence in the spinal cord neurons, followed by photoconversion using an UV light source (405 nm). Embryos were exposed to 30 flashes of 10 seconds UV light each, separated by 10 seconds of dark between flashes. After UV exposure, the embryos were immediately fixed in 4% PFA, followed by immunohistochemistry against FLAG (anti-FLAG, 1:200, Sigma, F1804) and EosFP Red (anti-CaMPARI-Red [4F6], 1:1000, Biozol, ABA-AB01649-23.0). Image acquisition was performed using the inverted Zeiss LSM 980 confocal microscope.

For analysis of neuronal activity, a square region of interest was drawn around all FLAG-positive somas in the spinal cord, followed by background subtraction and measurement of FLAG and EosFP Red mean intensity. Red/Flag ratio was used to assess the neuronal activity for each soma, and values were averaged per embryo. For quantification, the observer was blinded to the experimental condition.

### Behaviour analysis

To analyse the number of spontaneous contractions, 18 - 20 and 24 - 26 hpf not dechorionated embryos were recorded continuously for 5 min under a stereomicroscope (Zeiss Axio Zoom.V16 with a PlanNeoFluar Z 1.0x objective) equipped with an Axiocam 503 mono camera, and the number of spontaneous coiling movements counted (Oprişoreanu et al., 2019). For the experimental setup, embryos were maintained under constant ambient temperature and lighting conditions. Embryos were recorded individually, with one embryo imaged at a time. After transferring to a new dish, the single embryo was allowed to adapt for 1 min before recording. Genotypes were alternated during imaging (one control embryo followed by one *tac1* KO embryo). The videos were recorded using the Zeiss software at a resolution of 1936×1460 and speed of around 5 frames/second. Analyses of videos were done in a blinded fashion.

### Pharmacological inhibition of Tac1 receptor

To block the Tac1 receptor Tg(*mnx1*:GFP), eggs were incubated at 8 hpf with 50 µM of Rolapitant (APExBIO). The drug treatment was performed in 24-well plates, with 6 larvae per well. The stock concentration of the compounds was at 10 mM in DMSO and the final DMSO concentration in fish water was 0.5% DMSO. Vehicle controls were also at 0.5% DMSO. At 26 - 28 hpf, the embryos were manually dechorionated with insect pins and fixed in 4% PFA for 2 hours at RT. After fixation, embryos were washed with phosphate buffered saline with Tween-20 (PBST) 3 times for 5 min and stored in 70% glycerol in PBS. For CaP axonal morphology analysis, deyolked embryos were mounted onto glass slides and imaged using an inverse Imager microscope with an Apotome unit. CaP axon morphology scoring, total axon length, branch length and number of branches were analysed using the Fiji software.

## ACKNOWLEDGEMENTS

We thank Dr. Zhen Zhong for critically reading the manuscript; Dr. Judith Konantz, Marika Fischer, Silvio Kunadt and Denise Bärhold for fish care. The work was supported by the Light Microscopy Facility and the Zebrafish Facility, both core facilities of the Center for Molecular and Cellular Bioengineering (CMCB) at the Technische Universität (TU) Dresden. Funding: Alexander von Humboldt Stiftung Professorship Award (CGB) and TU Dresden Core Funding (CGB).

## AUTHOR CONTRIBUTIONS

AMO: conceptualization, methodology, analysis, investigation, validation, visualization, writing - original draft, supervision, writing - review and editing; SU: methodology, analysis, investigation; DZ: methodology, investigation; AB: investigation; TB: conceptualization, writing - original draft, visualization, supervision, funding acquisition, writing - review and editing; CGB: conceptualization, writing - original draft, visualization, supervision, funding acquisition, writing - review and editing.

## SUPPLEMENTARY FIGURES

**Suppl. Fig. 1:**
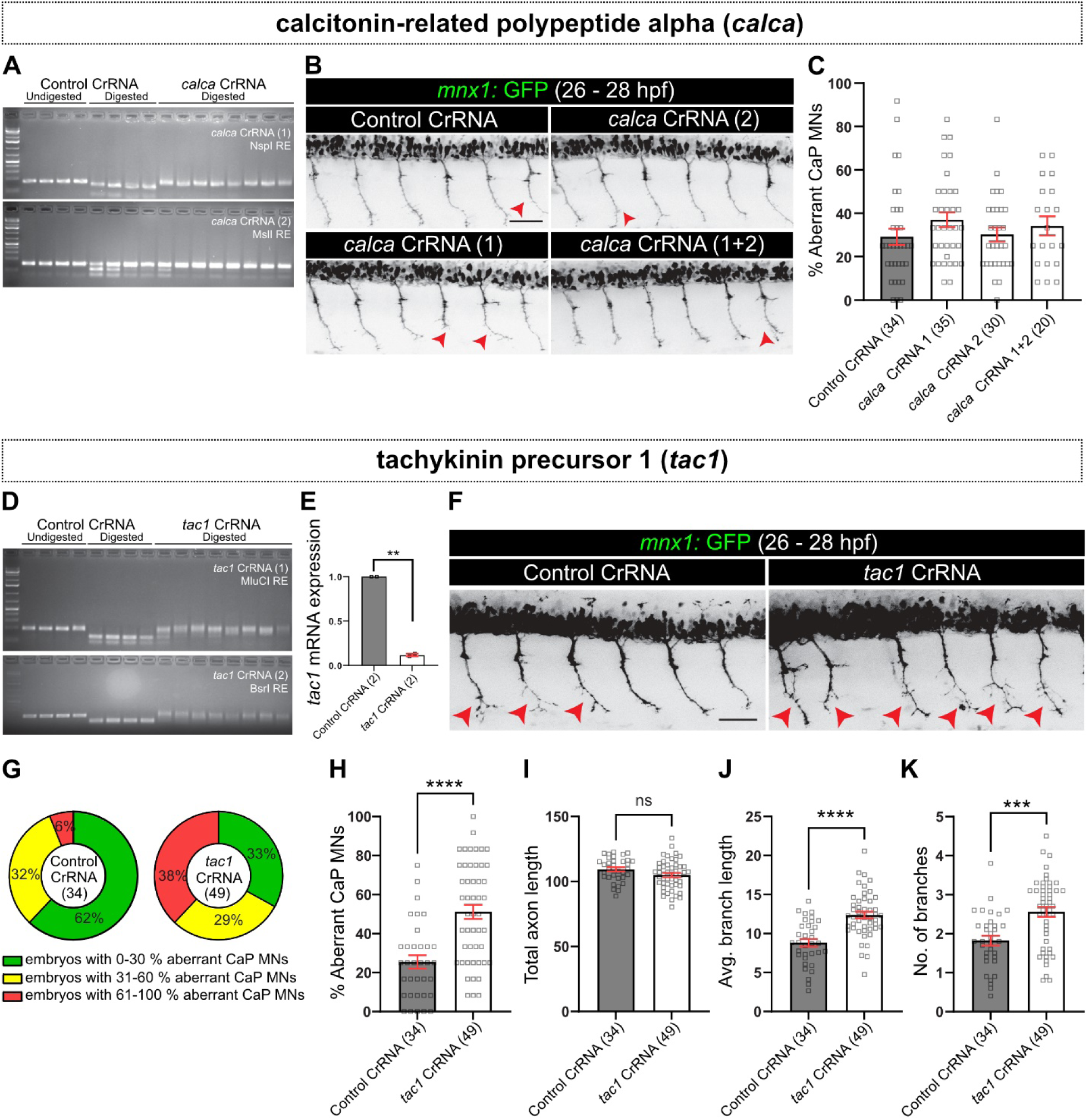
Functional screening of neuropeptide-coding genes in motor neurons indicates a role of *tac1* for CaP axon differentiation. **(A)** RFLP showing haCR gRNA injection efficiency for *calca*. Each lane represents one larva with and without digestion with NspI and MslI, respectively. **(B)** Representative images of 26 - 28 hpf *mnx1:GFP* transgenic embryos injected with *calca* and control CrRNAs. Red arrows indicate aberrant axons. **(C)** Quantification of CaP MNs show no aberrant phenotype in *calca* somatic mutants compared to control. **(D)** PCR and RFLP analysis used to determine the efficiency of the *tac1* CrRNA, as indel generation destroys the recognition site of the restriction enzyme. **(E)** qRT-PCR shows an 89% decrease in the *tac1* mRNA expression level in somatic mutants compared to control (one sample t-test, *tac1* CrRNA vs control CrRNA **p = 0.0100). **(F)** Lateral trunk view of 26- 28 hpf *mnx1*:GFP transgenic embryos. Red arrows indicate aberrant CaP axonal growth. **(G)** Distribution in percentage of the CaP axonal phenotype per experimental group. Green area represents the embryos that show between 0 and 30% aberrant CaP motor neurons, yellow area between 31 and 60%, and red area between 61 and 100%. **(H)** Quantification of aberrant CaP motor neurons (Mann Whitney test, ****p<0.0001). Analysis of the total axon length **(I)**, average branch length **(J)** and number of branches **(K)** in somatic mutants compared to control (avg. branch length, unpaired t-test, ****p<0.0001; no. of branches, unpaired t-test, ***p = 0.0002). Error bars show SEM. Scale bar: 50 µm.

**Suppl. Fig. 2:**
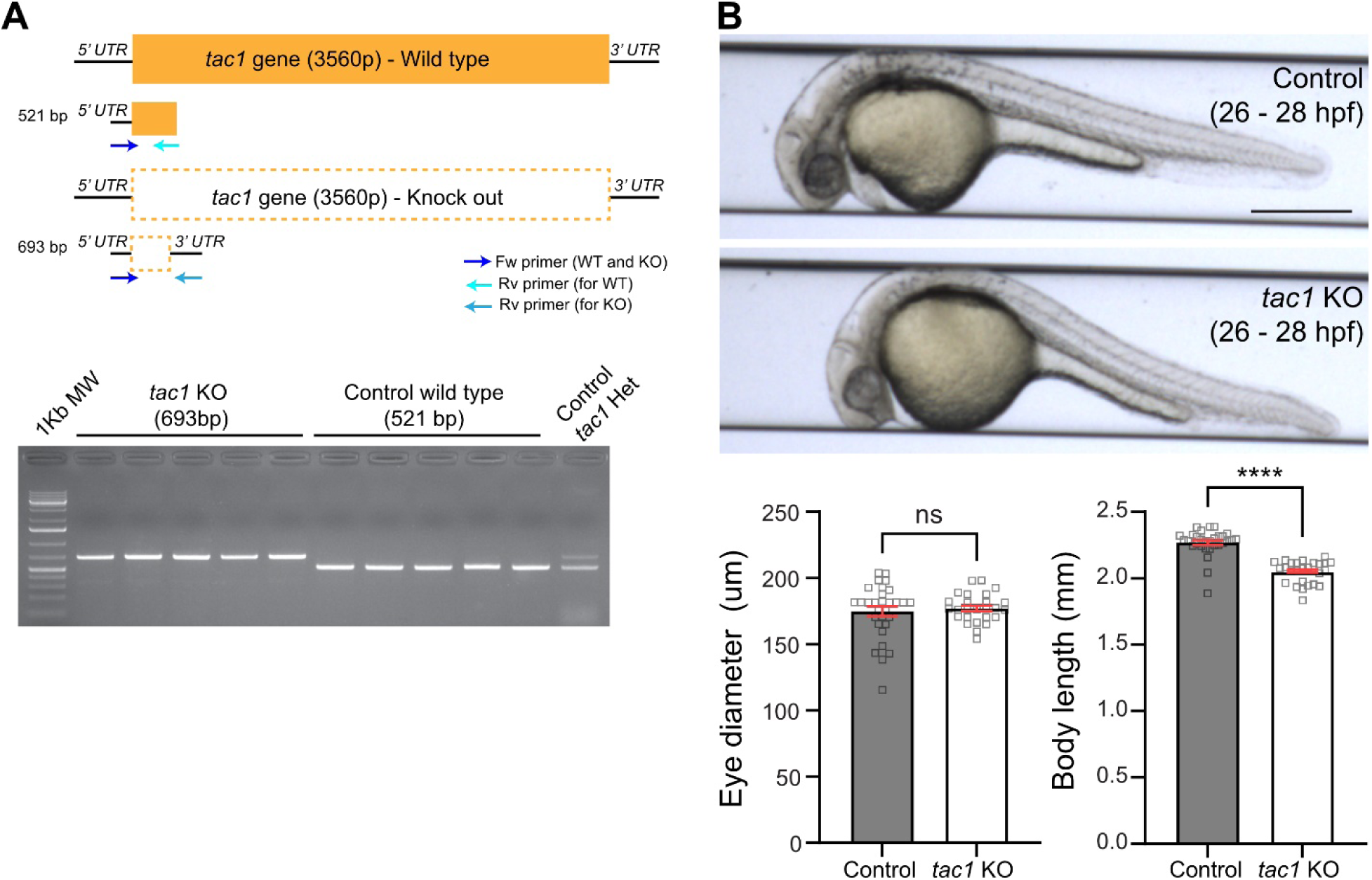
Generation of the *tac1* complete knockout. **(A)** Schematic drawing of the complete *tac1* ORF in wild-type and tac1 KO animals. The position of the genotyping primers is also noted. **(B)** VAST images showing wild-type and germline *tac1* KO embyos at 26 - 28 hpf. No significant changes in eye diameter are observed. *tac1* KO embryos display shorter bodies compared to wild-type control (Mann-Whitney, p**** < 0.0001). Scale bar: 500 µm. Error bars show SEM.

## Notes

### Competing Interest Statement

The authors have declared no competing interest.

